# Dissecting GPCR Selectivity: A complex interplay of various intracellular motifs determines G-protein binding and activation

**DOI:** 10.64898/2025.12.12.693856

**Authors:** Sina B. Kirchhofer, Volker Jelinek, Katharina Klingelhöfer, Anna-Lena Krett, Moritz Bünemann

**Affiliations:** Department of Pharmacology and Clinical Pharmacy, University of Marburg, Karl-von-Frisch-Str. 2, 35043 Marburg, Germany

## Abstract

G-Protein coupled receptors (GPCRs) mediate intracellular signaling by selectively activating heterotrimeric G-proteins. While certain GPCRs exhibit a high specificity toward particular G-protein subtypes, other GPCRs display promiscuous signaling by engaging interaction with multiple G-protein families. Molecular determinants underlying the selectivity or promiscuity of the receptors remain incompletely understood. In the present study, we investigate various structural motifs within the intracellular domains of Muscarinic receptors to assess their role in both Gα subunit binding and activation. To this end, we generated chimeric receptors and applied both FRET- and BRET-based assays to monitor G protein binding and activation. Our study demonstrates that the determination of G-protein coupling selectivity is not defined by single motifs or amino acids but rather by a complex interplay of various intracellular motifs affecting binding or/and subsequent activation. These results provide new insights into the structural basis of GPCR-G protein specificity.

## Introduction

G-Protein coupled receptors (GPCRs) are the largest family of transmembrane proteins, and through their contribution in countless physiological processes, they are one of the most important drug targets ^1–3^. GPCRs transmit extracellular signals via the heterotrimeric G-proteins (Gα, Gβ and Gγ). Activation of a GPCR triggers a conformational rearrangement within the receptor, facilitating the binding of heterotrimeric G-proteins at the cytoplasmic interface. This interaction promotes the exchange of GDP for GTP on the Gα-subunit, leading to activation and subsequent dissociation of the G-proteins ^3^. There are four different Gα-protein families (Gs, Gi/o, Gq/11 and G12/13) activating or deactivating various signaling cascades. Certain GPCRs exhibit a high selectivity through which G-protein they transmit their signal. In contrast, other GPCRs display promiscuous signaling profiles, coupling with Gα-subunits from one, two or even more different families. The mechanisms underlying this promiscuity in G-protein coupling, as well as the structural determinants within both the receptors and Gα subunits, have been the subject of extensive investigation over the past several decades. Although recent advances in crystal structures, Cryo-EM as well as molecular dynamics simulations have provided detailed structures of GPCRs in complex with the heterotrimeric G-proteins, the distinct motifs or conformational features that determine coupling selectivity remain unclear. Furthermore, it is still uncertain whether a universal coupling mechanism exists that applies across different GPCRs and G-protein families. However, some motifs have consistently been identified as key determinants of coupling specificity across different GPCR. These include the intracellular loops, the intracellular domains of the transmembrane helices 5 and 6, as well as the C-terminal domain. In this study, we engineered chimeric receptors incorporating the here named motifs from closely related GPCRs with distinct coupling selectivity. These chimeric receptors were subsequently analyzed for their ability to interact with different G-protein families via a Förster-resonance energy transfer (FRET)-based approach. For this, we employed a well-established biochemical assay in permeabilized cells under GTP depletion, which enabled further analysis of the binding stability of receptor-G-protein complex ^4,5^. In the next step, we assessed whether the G-proteins not only interact with the receptor, but also undergo subsequent activation. G-protein activation induced by the chimeric receptors was evaluated using the bioluminescence-resonance energy transfer (BRET)-based approach of the G-case sensors ^6^. These studies were conducted on members of the muscarinic acetylcholine receptor family, part of the aminergic receptors. In particular, we selected the muscarinic M_2_ receptor, which exclusively couples to Gi/o proteins, and the M_3_ receptor, which predominantly couples to Gq but also interacts with Gi/o in a less stable manner ^4^. We systematically analyzed the binding and subsequent activation of Gi/o and Gq-proteins by chimeric receptors incorporating motifs or combination of motifs of the respective other receptor. These analyzes provided new insights into a potential common mechanism underlying not only G-protein coupling selectivity, but also their subsequent activation, thereby influencing the induction of physiological responses.

## Results

### M_2_R-based chimeric receptors induce binding and activation of Gq proteins

To investigate the influence of intracellular domains of the M_2_ receptor (M_2_R) on G protein binding and activation, we reviewed previously published crystal structures, Cryo-EM structures, and other relevant studies to identify potential motifs crucial for receptor-G alpha subunit interaction. Using this information, we constructed chimeric receptors based on the M_2_R, incorporating specific motifs from the M_3_ receptor (M_3_R) (Fig. 1a, Supplemental Table 1) and vice versa. The motifs we identified as potentially relevant included the intracellular loop 2 and 3 (ICL2, ICL3), the intracellular domains of transmembrane helices 5 and 6 (TM56), the helix 8 (H8), a cluster of basic amino acids in the C-Terminus (polybasic cluster, pbc), the directly followed distal part of the C-Terminus (distal C-Term) and the complete C-Terminus (C-Term). These chimeric receptors were subsequently characterized for their ability to interact with G alpha o or G alpha q proteins. To this end, we used a FRET-based approach under GTP-depletion in HEK293T cells transfected with the receptor or chimeric receptor with a C-Terminal mCitrine, Gα_q_ or Gα_o_, Gβ_1_, and CFP-Gγ_2_ (Fig. 1b) and performed single-cell measurements as described before ^4^. We further transfected a plasmid coding for PTX-A, to inhibit any effects transmitted via endogenous G alpha i/o proteins, whereas the transfected Gαo proteins were rendered to be PTX insensitive due to a point mutation in the C-terminus. For the measurement, cells were permeabilized by exposure to 0.05% saponin for 2 min, depleting the membranes from nucleotides and increasing the stability of the agonist-bound receptor – G-protein complex. During the measurement, cells were superfused with buffer, agonist or GTPγs solution. The dissociation kinetics of this ternary complex after wash-out of the agonist acetylcholine were quantified by the calculation of the area under the curve (AUC) as described before ^5^ (Figure 1c-e, Supplemental Figure 1). The complete dissociation of the complex was induced by the application of GTPγs. The wildtype M_2_R only interacted with Go proteins, whereas the wildtype M_3_R was able to interact with both Go and Gq proteins, generating a more stable complex between M_3_R and Gq than with Go (Fig. 1e; as shown in ^4^). To analyze if the chimeric receptors were not only able to interact but also subsequently activate the G proteins, we applied a BRET-based approach using the activity sensors Go_1_-case or Gq-case ^6^ as described before ^7,8^ (Figure 1f). For this, cells were transfected with the receptor chimeras and the respective activity sensor. The receptor chimeras were fused with mCherry at the N-terminal and comparable membrane expression was verified via flow cytometry (Supplemental Figure 2). For the measurement, we applied an increasing concentration of carbachol (CCh), followed by a fully saturating concentration of 300 µM CCh for normalization. For the multiwell-based BRET measurements we used CCh to activate the muscarinic receptors as it has a higher stability and did not need to be washed off the receptor. We fitted concentration-response curves (Fig. 1g, Supplemental Figure 2) and calculated the EC_50_-values (Fig. 1h). The M_2_R induced a sufficient activation of Go (EC_50_: 480 nM), but not of Gq, whereas M_3_R was able to activate both Go (EC_50_: 1 µM) and Gq (EC_50_: 170 nM), with a higher affinity for Gq (Fig. 1f-g).

**Figure 1:**
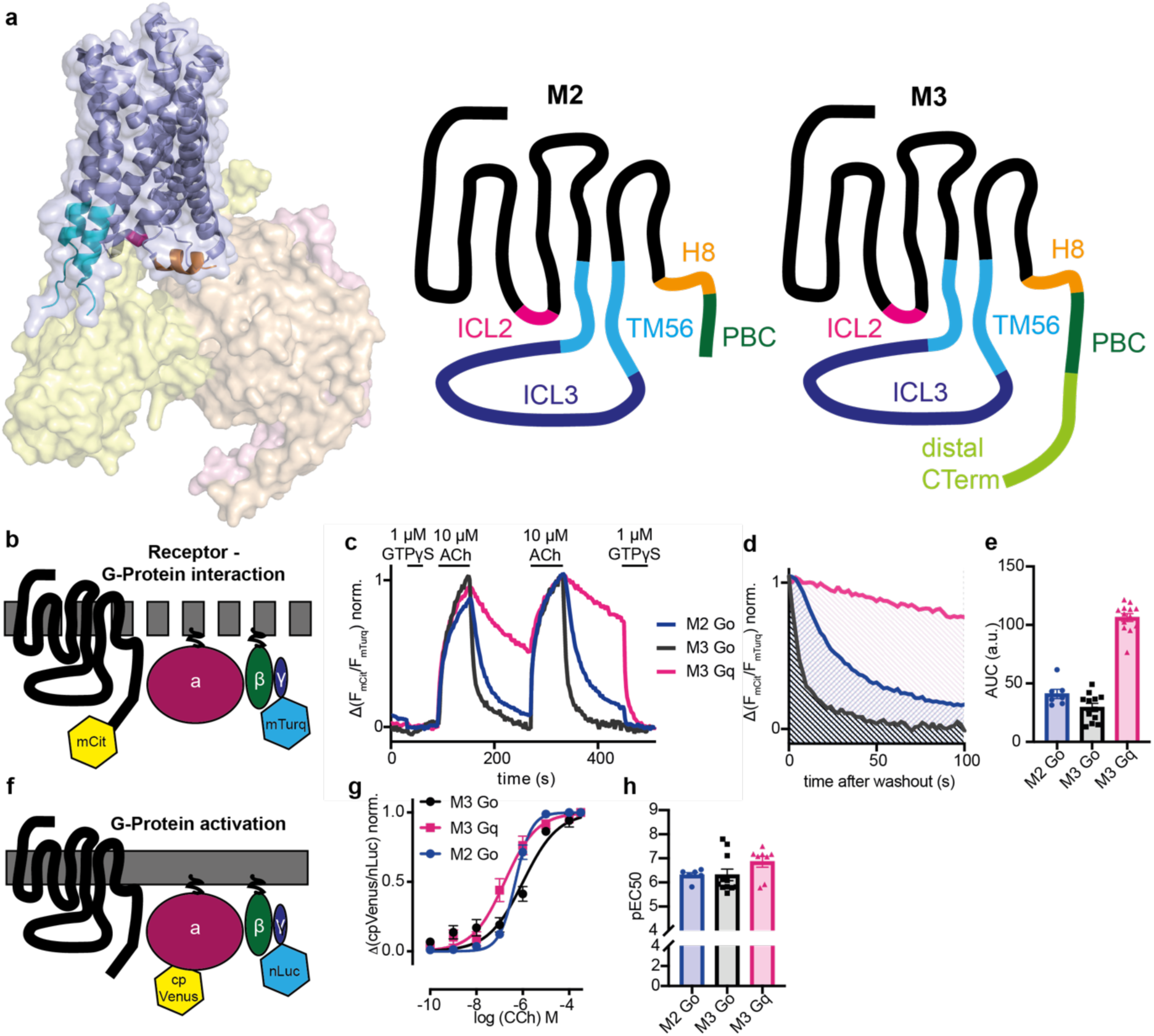
Chimeric Muscarinic Receptors allow the measurement of Receptor – G-Protein Interaction and G-Protein activation. **a** Scheme of the M_2_-receptor in complex with the heterotrimeric G-proteins (Gαi (yellow), Gβ (orange) and Gγ (pink)) (based on PDB: 7T8X). Different intracellular parts were exchanged between M_2_R and M_3_R, depicted in the comic. **b** Scheme of the performed FRET-based assays for the measurement of receptor-G-protein interaction performed in permeabilized HEK293T cells expressing a mCit-tagged receptor and the respective G-proteins with a Gγ carrying a mTurq. **c-e** Representative FRET-based measurement of receptor-G-protein interaction in transfected, permeabilized HEK293T cells. Cells were superfused with the depicted solutions or buffer. The area under the curve (d) for the time after wash out of the second 10 µM acetylcholine application was calculated to quantify the strength of the interaction and plotted as bar graph (e). **f** Scheme of the performed BRET-based assays for the measurement of G-protein activation in HEK293T cells expressing the G-case assays ^6^. **g-h** G-protein activation was measured BRET-based in multiwell format in a plate reader and concentration response curves for the carbachol induced G-protein activation were fitted (g). The calculated EC50-values were plotted in a bar graph (h).

Chimeric M_2_ receptor constructs were analyzed as described in Figure 1. The chimeric M_2_R carrying TM56 and Helix 8 of M_3_R did not display any interaction with Go proteins (Fig. 2a-b), but showed a stable interaction with Gq. Same was true for chimeric receptors with the TM56 or TM56 combined with ICL3 of M_3_R (Figure 2b). These chimeric receptors were further able to subsequently activate Gq proteins, but not Go proteins (Figure 2c-d), changing the selectivity of the M_2_R from explicit Go coupling to explicit Gq. Chimeric M_2_R carrying the TM56 in combination with other motifs (pbc, ICL3+pbc or ICL3+H8+CTerm) were able to bind both Go and Gq proteins (Figure 2e), but only induced activation of Gq (Figure 2f). Only the exchange of ICL2 in addition to ICL3 and TM56 resulted in chimeric M_2_R that was able to bind Go and Gq in comparable manner (Figure 2g) and furthermore activated both G proteins (Fig. 2h). Overall, the exchange of the intracellular domains of the helices 5 and 6 from M_3_R into M_2_R notably altered the selectivity of the M_2_R, shifting its coupling and activation profile from Go to Gq. Combinations of these regions with the C-Terminus, or isolated parts of the C-Terminus, rescued Go binding, although activation was not observed. Finally, the additional incorporation of ICL2 further enabled Go binding and subsequent activation.

**Figure 2:**
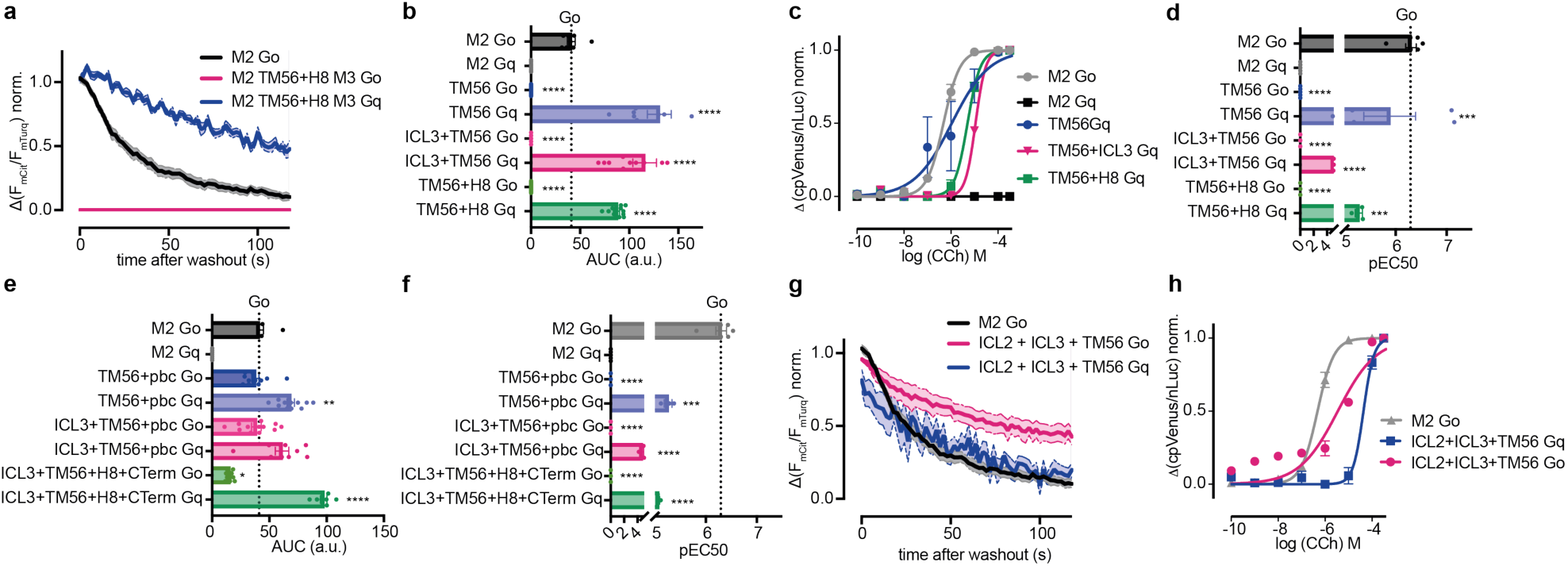
M_2_R-based chimeric receptors enable the binding and subsequent activation of Gq proteins by the M_2_ receptor. **a-b** Averaged measurement of receptor Go or Gq-protein interaction of M_2_R and chimeric M_2_R with TM56 and Helix 8 of M_3_R. The chimeric receptors with TM56, TM56 combined with ICL3 or TM56 and Helix 8 of M_3_R did not interact with Go, but enabled interaction with Gq. The calculated AUC is blotted as bar graph in b. **c-d** Concentration-response curves for chimeric M_2_/M_3_ receptors induced G-protein activation (c), calculated EC50 values were plotted in bar graph (d). **e-f** M_2_/M_3_ chimeras with TM56 combined with pbc, ICL3 and pbc or ICl3, Helix 8 and C Terminus of M_3_R enabled the binding of Go and Gq (e) but only subsequently activate Gq proteins (f). **g-h** Chimeric receptor of M_2_R with ICl2, ICL3 and TM56 from M_3_R allowed the interaction (g) and activation (h) of both Go and Gq proteins. (Statistic: Ordinary One-way ANOVA with Dunnett’s multiple comparison test, **** p<0.0001, *** p<0.001, ** p<0.01, ns: p>0.05).

### M_3_R-based chimeric receptors can signal solely via Gq or Go

Having successfully altered the G-protein coupling selectivity of the M_2_R, we applied the same approach on the M_3_R (Figure 3, Supplemental Figure 3-4). The M_3_R can signal through both Gq and Go proteins, however, favors the Gq pathway (Gq: EC_50_: 170 nM, AUC: 106 a.u; Go: EC_50_: 1 µM, AUC: 30 a.u.). The insertion of the TM56 and H8 motif from M_2_R into M_3_R abolished the binding (Figure 3a) and activation (Figure 3b) of Gq and instead, increased binding and activation of Go (Figure 3a-b). The motifs ICL3 and TM56 allowed a highly stable interaction of Go with the receptor, but were unable to induce a detectable interaction with Gq (Fig. 3c). However, this chimeric not only strongly activated Go, but at higher agonist concentrations also induced Gq activation (EC_50_: Gq: 1,9 µM vs. Go: 35 nM, Figure 3d). Similar effects were observed for the combination of TM56 and pbc (Figure 3e-f), indicating that both chimeras bind and activate Go. However, these two chimeras appear to interact weekly or transiently with Gq, although this interaction is still sufficient for activation. The full exchange of all above mentioned motifs from M_2_R into M_3_R, gave rise to functional receptors that strongly activated Go (EC_50_: 30 nM), without any Gq activation. However, fusing this chimeric receptor to a C-terminal mCitrine prevented proper expression at the plasma membrane. Therefore, we were unable to study G-protein binding by means of FRET. The chimera without the c-terminal mCitrine expressed comparable to WT as detected by fluorescence of receptors N-terminal fused mCherry-version (Supplemental Figure 4). The exchange of the pbc together with truncation of the C-Terminus after the pbc facilitated the binding (Figure 3g) and activation (Figure 3h) of solely Gq. Truncation of C-Terminus alone, as well as the complete exchange of helix 8 and C-Terminus together resulted in receptors where only Gq binding was observed (Figure 3i), however, activation of both Gq and Go could be detected (Figure 3j). This indicates a comparable behavior as shown for the chimeric M_3_R with TM56 and pbc or ICL3 for Gq activation without detectable binding to the receptor (Figure 3e-f). The M3-based chimeric receptor with ICL2 and 3 and TM56 allowed the binding of both Go and Gq (Figure 2k), but activated only Go and not Gq (Figure 3l), with a remarkably increased coupling efficacy for Go activation (EC_50_: 2nM). The additional insertion of ICL2 seems to rescue the stability of Gq-receptor complex (Figure 3c and k), but diminishes its activation (Figure 3d and l). All in all, as for M_2_R, the switch of the intracellular parts of the helices 5 and 6 with combinations of other motifs from M_2_R into M_3_R changed the coupling-selectivity for M_3_R to favoring Go. Modifications in the C-Terminal region of the receptor lead to a switch in coupling-selectivity, favoring Gq. However, several motif combinations reduced the stability of G-protein-receptor complex to levels undetectable with our methods, yet still allowed the subsequent activation of both G-protein variants.

**Figure 3:**
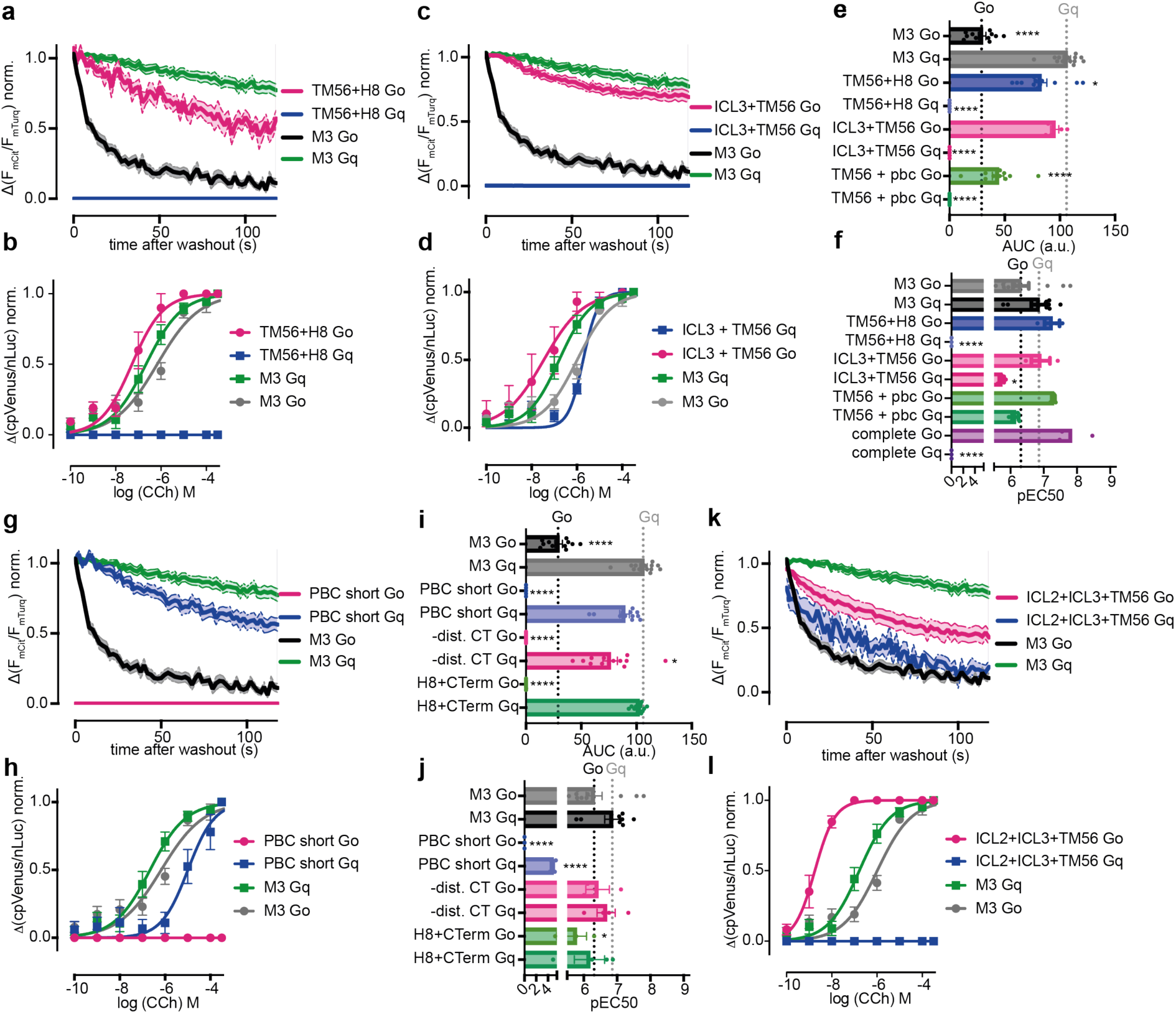
M_3_R-based chimeric receptors can be modified to signal solely via Gq or Go. **a-b** Averaged measurement of receptor Go or Gq-protein interaction of M_3_R and chimeric M_3_R with TM56 and Helix 8 of M2 (a). The chimeric receptor only interacts with Gq. Concentration response curve for the G protein activation (b). The M_3_R TM56 and Helix 8 chimera only activated Go proteins. **c-d** Averaged measurement of receptor Go or Gq-protein interaction of M_3_R and chimeric M_3_R with TM56 and ICL3 of M_2_R (c). The chimeric receptor only interacts with Gq. Concentration response curve for the G protein activation (d). The M_3_R TM56 and Helix 8 chimera is able to activate both Go ang Gq. **e-f** Bar graphs of the calculated area under the curve for chimeric M_3_R with TM56 of M_2_R combined with helix 8, ICL3 or PBC (e). Calculated pEC50 values plotted as bar graphs (f). Chimeric M_3_R with TM56 and Helix 8 or a complete exchange to M_2_R only activated Go, the other chimeras also activated Gq. **g-h** Receptor-G-protein interaction between M_3_R with the PBC of M_2_R and a truncated C-Terminus (g). The chimera only interacted with Gq. The chimera as well only activated Gq (h). **i-j** Bar graphs of the calculated area under the curve (i) for chimeric M_3_R with modulated C-terminus. Calculated pEC50 values plotted as bar graphs (j). For chimeric receptors with the modulated C-terminus no interaction with Go was detectable. Chimerics still carrying the M_3_R PBC could still activate Go. **k-l** Receptor-G-protein interaction between M_3_R with ICL2, ICL3 and TM56 of M_2_R. The chimera displayed an increased interaction with Go and a decreased interaction with Gq (k) and only activated Go (l). (Statistic: Ordinary One-way ANOVA with Dunnett’s multiple comparison test, **** p<0.0001, *** p<0.001, ** p<0.01, ns: p>0.05).

### Various motifs define the coupling selectivity of M_2_ and M_3_ receptors

We examined all different chimeric receptors between M_2_R and M_3_R with various combinations of motifs, for their binding and activation profiles with respect to Go vs. Gq (Figure 4). Overall, combinations with TM56-motif decreased Go binding and abolished activation by M_2_R (Figure 4a and d) but increased it for M_3_R (Figure 4e and g). Additionally, these chimeras allowed Gq binding and activation by M_2_R (Figure 4b and d), which was decreased for M_3_R (Figure 4f and h). Modulations of just the C-Terminal part of the receptors led to a decrease in Go binding of M_3_R (Figure 4e) and an increase for M_2_R (Figure 4a). Overall, we were able to create M_2_R-based chimeric receptors which were now explicitly Gq-coupled receptors. Further, we created M_3_R-based chimeric receptors which were now selective for one G-protein, Go or Gq. Moreover, we demonstrated that the binding of a G-protein to a receptor does not necessarily result in activation. It appears that different motifs of the receptor are responsible for binding, while others contribute to activation through their interactions. Additionally, some motifs, such as ICL2 or 3, have a minimal effect on their own. However, in combination with other motifs, they can restore a previously diminished interaction.

**Figure 4:**
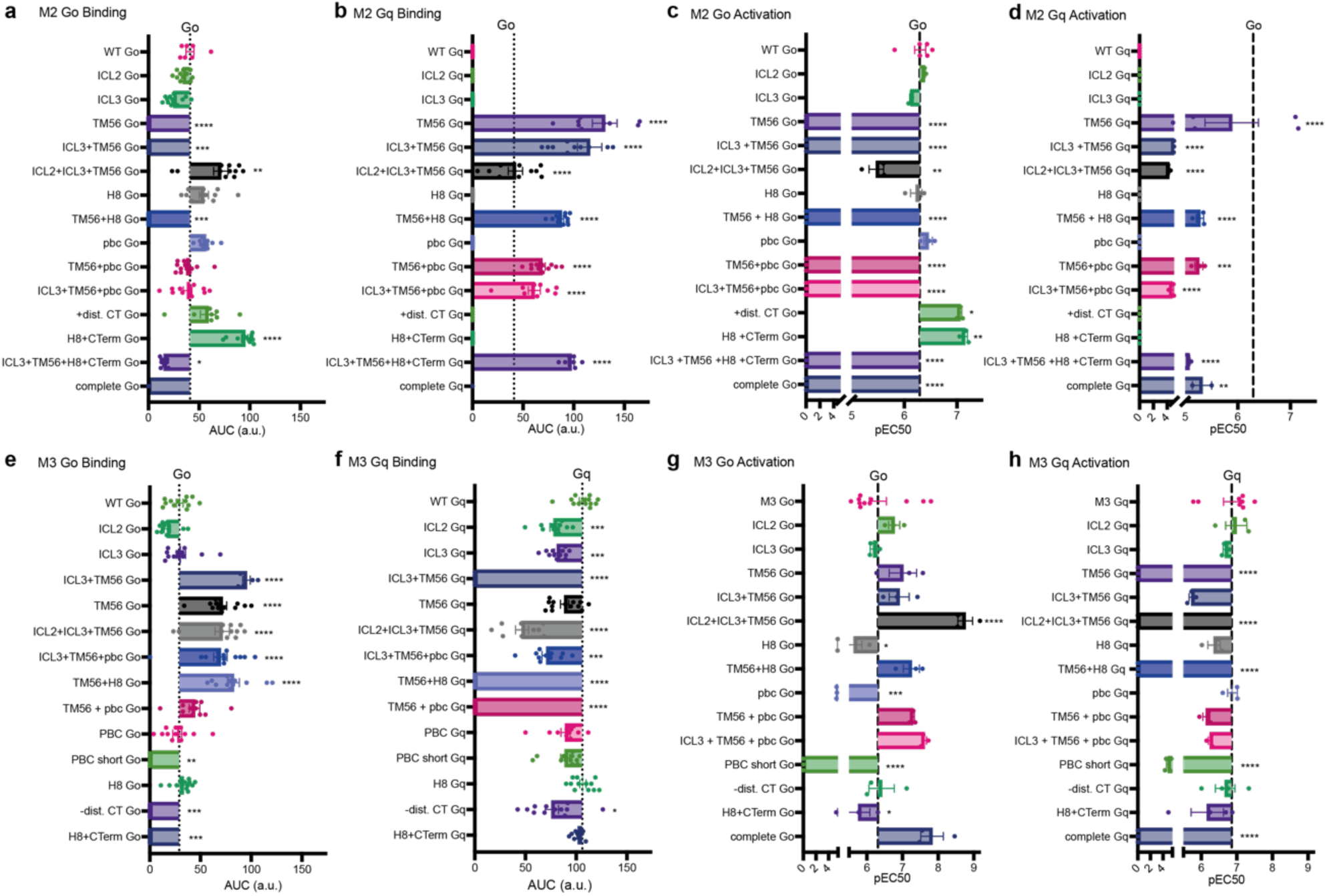
Chimeric receptors between M_2_ and M_3_ receptor modify both interaction and activation of Go and Gq proteins. **a-d** Bar graph showing the effects on Go binding (a), Gq binding (b), Go activation (c) and Gq activation (d) of all M_2_R-based chimerics in comparison to the Go binding or activation induced by the WT receptor. **e-f** Bar graph showing the effects on Go binding (a), Gq binding (b), Go activation (c) and Gq activation (d) of all M_3_R-based chimerics in comparison to the G o or Gq binding or activation induced by the WT receptor.

Taken together, Go coupled receptors exhibited enhanced affinity to Go upon the insertion of C-terminal regions of Gq/o coupled receptors, while the insertion of intracellular parts of transmembrane helices 5 and 6 reduced their Go-affinity (Figure 5). This switch facilitated Gq binding and signaling in Go coupled receptor (shown for Go, Figure 2,4,5b). Gq/o coupled receptors exhibited reduced affinity to Go by insertion of C-terminal parts of Go coupled receptors, while their Go-binding and activation increased by incorporation of intracellular parts of transmembrane helices 5 and 6 (shown for M_3_R, Figure 3,4,5c), which also led to a decrease of the the binding and especially activation for Gq (shown for M_3_R, Figure 3,4).

**Figure 5:**
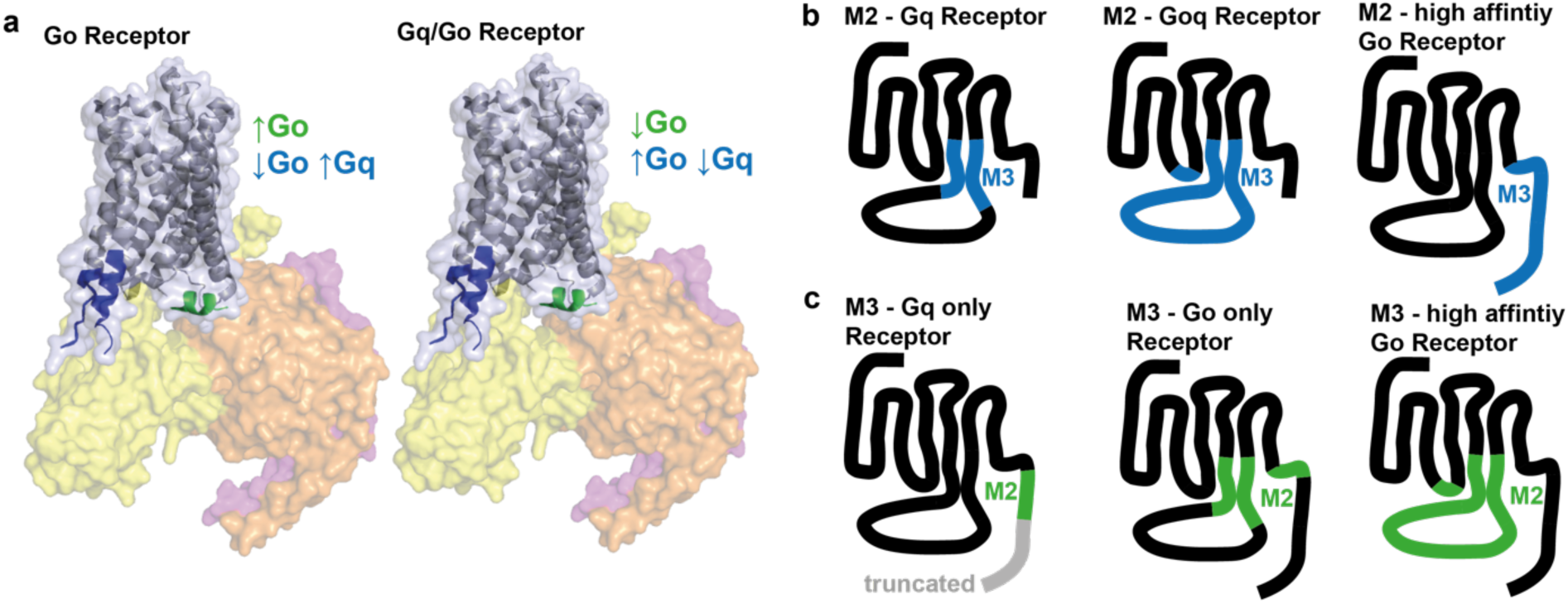
Different intracellular motifs in Class A GPCRs influence the ability of a receptor to interact and subsequently activate Go or Gq proteins. **a** Scheme of a receptor in complex with the heterotrimeric G-proteins (Gα (yellow), Gβ (orange) and Gγ (pink)) (based on PDB: 7T8X). Go coupled receptors increase their Go-affinity by insertion of C-terminal parts of Gq/o coupled receptors (green) and decrease their Go-affinity by insertion of intracellular parts of transmembrane helices 5 and 6 (blue), which also enables Gq binding. Gq/o coupled receptors decrease their Go-affinity by insertion of C-terminal parts of Go coupled receptors (green) and increase their Go-affinity by insertion of intracellular parts of transmembrane helices 5 and 6 (blue), which also decreases Gq-affinity. **b** Scheme of different M_2_R-based chimeric receptors (black) with motifs from M_3_R (blue). M_2_R is acting as Gq receptor when carrying the TM56 of M_3_R, it couples and activates both Go and Gq when carrying ICl2, ICL3 and TM56 of M_3_R. Change of H8 and C-Terminus increases the affinity for Go activation. **c** Scheme of different M_3_R-based chimeric receptors (black) with motifs from M_2_R (green). The M_3_R couples only Gq when carrying the pbc of M_2_R and a truncated C-Terminus. It couples only Go with the TM56 and H8 of M_2_R. Change of ICL2, ICL3 and TM56 increases the affinity for Go activation.

## Discussion

The molecular basis underlying G-protein coupling selectivity remains an area of intensive research. Numerous studies have explored potential important motifs within both G-proteins and GPCR itself, however, a universal mechanism has yet to be established. In this study, we systematically examined the functional effects of previously proposed motifs within the intracellular parts of GPCRs exhibiting divergent G-protein coupling selectivity. To this end, we engineered chimeric receptors by combining muscarinic M_2_ and M_3_ receptors. These chimeric receptors were analyzed by FRET- and BRET-based assays to assess their ability to first interact with the specific G-protein subtype, and second subsequent activate it.

All in all, our results demonstrate the ability to alter the G-protein coupling selectivity of different receptors, effectively converting receptors from selective to promiscuous couplers and vice versa. Through chimeric receptor analysis, we identified the cytoplasmic end of transmembrane helices 5 and 6, the ICL3 and parts of the C-terminal tail as the intracellular regions that most significantly determine coupling specificity. These regions have previously been implicated as key determinants of G-protein selectivity, particularly in the context of the “bar code” hypothesis ^9^ and have been supported by other, mainly computational, studies ^10–14^. However, especially the C-terminal part of GPCRs as well as the large ICL3 of muscarinic receptors are mainly not resolved in the crystal or Cryo-EM Structures of GPCR-G-Protein-complexes ^15–21^, making the computational evaluation of the important residues difficult. The functional relevance of larger intracellular motifs is frequently investigated by the use of modified or chimeric receptors, as employed in the present study. Consistent with previous reports, no universal motif responsible for coupling specificity could be found. For example, we observed that the truncation of the C-terminus reduced the coupling of Go and Gq to M_3_, but modulations of the ICL3 alone displayed only minor effects, especially did not affect the G-protein activation – confirming the observations of Ham and colleagues ^14^ as well as Inoue and colleagues ^10^. Moreover, we could show that the incorporation of intracellular domains of TM5 and 6 along with ICL3, promotes preferential coupling to Go proteins, which is as well in agreement with Inoue et al ^10^.

Looking on the ICL2, our data suggest no significant role for this intracellular loop on G-protein selectivity in isolation. This is contrary to some studies which have proposed ICL2 as one of the key determinants of G-protein selectivity ^22^. Based on our data, its influence appears to be context-dependent, emerging only in combination with other modifications. Additionally, previous findings indicate that the ICL2 predominantly interacts with Gs-proteins rather than Gi/o ^23^, and is not critical for receptors that primarily couple to Gi/o, such as the M_2_ receptor ^24^. This aligns with our own observations. Furthermore, other studies based on the class B GPCR PTH1 showed that Gq signaling is even more depended on the ICL2 than Gs signaling ^25^. Kim and colleges ^24^ further noted that the ICL2 may play a more prominent role when Gi/o is a secondary coupling partner, where its interactions with the G-protein binding pocket are comparatively weaker. This may explain our observations upon modulation of TM56 and ICL3 in the M_3_ receptor, where only modest changes in Go activation were detected. However, subsequent incorporation of the ICL2 from the M_2_ receptor strongly enhanced Go activation by the M3 receptor. In line with the majority of our findings, the most pronounced effects on G-protein selectivity were observed when multiple intracellular motifs were modified in combination, rather than individually. This underscores the cooperative nature of the structural determinants involved in coupling specificity. Previous studies have demonstrated that selectivity of G-protein binding involves multiple types of receptor-G-protein contacts, and not only several different contacts are necessary, but also the temporal life time of the complex is critical for effective signaling ^4,12^. Further, not single residues seem to be the determinant, selectivity appears to be influenced by broader structural features such as the size of the G-protein binding pocket, the insertion site ^21,26^, the total number of receptor-G-protein contacts ^27^, the conformational state of the receptor ^23^ and the angle for insertion of G-Protein into the receptor core ^28^. It is well established that Gs proteins require a comparatively wider G-protein binding pocket, whereas Gi/o-proteins can engage more compact binding cavities ^17,21^. Additionally, GPCRs that couple solely to Gi/o are generally more selective, while Gq-coupled GPCRs often exhibit at least some degree of promiscuous coupling to Gi/o proteins ^26^. Furthermore, similar to Gs proteins, Gq proteins appear to require a wider opening of the G-Protein binding pocket ^23^. It has also been reported that Gi/o require fewer receptor interaction contacts compared to other G-protein subtypes ^27^. Notably, the M_3_ receptor is known primarily couple to Gq, while also exhibiting weaker, less stable interactions with Gi/o, as well as with Gs and G13 ^29^, suggesting an already structural flexible or diverse binding pocket. In this study, we assessed not only the agonist-induced binding probabilities of receptor and G-proteins, but also the subsequent agonist-induced activation of G-proteins downstream. In some chimeric receptors, we observed stable G-protein binding without efficient activation, indicating that receptor-G-protein association alone is not sufficient to drive further signal transduction. As said before, it was shown that the conformational state of the receptor is influencing the coupling selectivity of the GPCR ^23^, indicating that the agonist induced conformational change induces the binding of the G protein, without directly activating it, also shown for some of the chimeric receptors. Further studies also investigated this effect. Morales-Pastor and colleagues ^30^ were able to show that the allosteric communication network within the receptor has an impact on the protein function, and with this the coupling selectivity of the GPCR. This demonstrates that the coupling selectivity of GPCRs already starts with the ligand binding, continuing with conformational rearrangements within the receptor, the binding of the G proteins and finally their subsequent activation. Comparable effects can also be seen in our study. Modulations of the intracellular parts of TM56, parts of the G-protein activating domain ^15,31^, still allow the binding of Go and Gq to the M_3_ receptor, but the Go activation is strongly increased whereas the Gq activation is diminished. The opposite effect can be seen for the M_2_ receptor, where the activation of Go is prohibited, but activation of Gq is allowed. Furthermore, we made the surprising observation that some receptor chimeras were able to activate G proteins, however no agonist-induced binding was detectable (Fig. 3 I,j) indicating that G protein activation does not necessarily require a high stability of the ternary complex. Comparable effects were observed by Ham and colleagues when modulating the positively charged amino acids in the C-terminus of the M_3_ ^14^. This underscores the importance of evaluating both binding and activation events when investigating determinants of G-protein selectivity. As such, studies like ours are essential to move beyond structural or computational predictions alone, emphasizing the value of integrating multiple functional assays to comprehensively characterize GPCR-G-protein interactions.

Taken together, our findings demonstrate that G-protein coupling selectivity is not determined by a single motif or individual amino acid residue. Rather, it emerges from the complex interplay of multiple intracellular structural elements. We show that selective modulation of these regions – particularly in combination – can reprogram the coupling specificity of GPCRs. For example, we successfully redirected the highly selective M_2_ receptor toward an alternative signaling pathway, and conversely, transformed the broadly coupling M_3_ receptor into a more selective one. These results underscore the modular and dynamic nature of GPCR–G-protein interactions.

Collectively, these findings support the conclusion that various intracellular motifs within GPCRs critically influence first the selectivity to bind a specific G-protein and second the receptor’s ability to effectively subsequently activate the bound G-protein.

## Materials and Methods

### Plasmids

cDNAs for the human M_3_-mCit receptor ^5^, PTX-insensitive Gαo ^32^, Gαq ^33^, Gβ_1_-wt, mTurq-Gγ2 ^34^ and BRET-based activity sensor Go_1_-case and Gq-case ^6^ have been described before. PTX-A was purchased from Addgene (catalogue number 16678, Addgene, Watertown, Massachusetts, USA) and cloned into pcDNA3 expression vector. Si-mCherry-M_3_ receptor was constructed based on M_3_-mCit and mCherry ^35^. Si-mCherry-M_2_ and M_2_-mCit receptors were constructed based on the human M_2_ receptor ^36^. All constructed receptor chimeras can be found in Supplemental table 1, describing all exchanged positions. All oligonucleotide primers used in this study were designed using SnapGene Viewer (GSL Biotech, San Diego, California, USA), purchased at Eurofins Genomics (Ebersberg, Germany) and are listed in Supplemental Table 2. Plasmids were constructed using the NEBuilder Hifi DNA assembly kit or the Q5-site directed mutagenesis kit from New England Biolabs (Ipswich, USA).

### Reagents

DMEM, FCS, PBS with or without Ca^2+^ and Mg^2+^, penicillin/streptomycin, L-glutamine and trypsin-EDTA were purchased from Capricorn Scientific (Ebsdorfergrund, Germany). BSA (protease-free), Carbachol, Acetylcholine iodide and GTPγS were purchased from Sigma-Aldrich (St. Louis, Missouri, USA). Saponin was purchased at AppliChem (Darmstadt, Germany). Protease inhibitor cocktail tablets c0mplete mini EDTA-free were purchased at Roche (Basel, Switzerland). PEI (polyethylenimine, linear) was purchased at Polysciences (Warrington, Pennsylvania, USA). 6H-F-Colenterazine was provided by Dr. Wibke Diederich (University of Marburg, Germany) and was used as published previously ^7,8^.

### Cell culture

All experiments in this study were carried out in HEK293T, which were a kind gift from the Lohse laboratory, University of Würzburg, Germany. Cells were cultured in DMEM (Dulbecco’s Modified Eagle’s Medium, 4.5 g L-1 glucose) supplemented with 10 % FCS, 2 mM L-glutamine, 100 U/ml penicillin and 0.1 mg/ml streptomycin at 37°C and 5% CO_2_ and were tested for mycoplasma on a regular basis. For FRET measurements, HEK293T cells were transiently transfected with 0.8 µg of the respective receptor-construct labeled with mCit, 1.5 µg Gαq or Gαo, 0.5 µg Gβ_1_-wt, 0.2 µg mTurq-Gγ2 and 0.3 µg PTX-A in 6 cm Ø dishes using Effectene Transfection Reagent according to manufacturer’s instructions (Qiagen, Hilden, Germany) two days before the measurement. For BRET-based measurements, HEK293T cells were transiently transfected two days before the measurement using PEI reagent. For the measurement of G-Protein activity, cells were transiently transfected with 1.5 µg of the respective receptor construct and 1.5 µg of the BRET-activity sensor Go1-case or Gq-case (100 ng DNA/well). The mixing ratio of PEI to DNA was 3:1 with 1 mg/ml PEI. Per 1 µg DNA, 50 µl serum-free DMEM was added. The DNA-DMEM and PEI-DMEM mix were incubated for 5 minutes, then they were combined and further incubated for 10 minutes. The cells were counted and set to 300.000 cells per ml (30.000 cells / well), the DNA-PEI mix was added and seeded onto white sterile poly-L-lysine coated 96-well microplates and incubated for 48h. For expression control of the chimeric receptors, HEK293T cells were transiently transfected with 0.75 µg DNA using PEI as described above. 60.000 cells / well were seeded onto 24-well plates and incubated for 48h. Again, cells transfected with purinergic receptors were incubated with the respective antagonist during cell culture.

### Single-cell FRET measurements in permeabilized cells

Single-cell FRET measurements were performed at an inverted microscope as described before ^5^. The measurements were carried out at room temperature at an inverted microscope (Axiovert 100, Zeiss, Oberkochen, Germany), equipped with a 60x oil-immersion objective (UPlanSApo x60/ 1,35 Oil, Olympus, Tokio, Japan), LED light source with 440 nm and 500 nm (pE-100, CoolLED, Andover, United Kingdom) and a sCMOS-camera pco.panda 4.2 (Excelitas Technologies, Waltham, Massachusetts, USA). During the measurement, CFP was excited using an excitation filter (436/20, Chroma Technology, Bellows Falls, Vermont, USA) and a dichroic beam splitter (458LP, Semrock, Rochester, New York, USA). Fluorescence emission was collected simultaneously for YFP and CFP by a beamsplitter (505LP, Chroma) and two emission filters (CFP: 470/24, Chroma and YFP: 525/39, Semrock). Cells were illuminated by 60ms light flashes with a frequency of 0.5 Hz. During the measurement, cells were superfused using a pressure-driven perfusion system (VC3-8xP, ALA Scientific Instruments, Farmingdale, New York, USA). Data were collected by the VisiView software (Visitron Systems, Puchheim, Germany). For analysis of the receptor – G Protein interaction, cells were permeabilized as described before ^4^. For this, transfected cells were sed one day prior the measurement on Poly-L-Lysin coated coverslips. Prior the measurement, the coverslip with cells was fixed in a microscope chamber and washed with external buffer (NaCL 137 mM, KC 5.4 mM, HEPES 10 mM, CaCl_2_ 2 mM, MgCl_2_ 1 mM, pH 7.35). To permeabilize the cells, coverslips were incubated for 2 min in internal buffer (K^+^ aspartate 100 mM, KCl 30 mM, NaCl 10 mM, HEPES 10 mM, EGTA 5 mM, MgCl_2_ 1 mM, pH 7.35) containing 0.05% saponin. Afterwards, the coverslip was washed 5 times with internal buffer and single cells were measured as described before ^4,5^. During the measurement, cells were superfused with internal buffer or internal buffer containing 10 µM Acetylcholine or 1 µM GTPγS using a protocol shown in Figure 1c.

### BRET measurements

BRET measurements were performed using a Spark 20M plate reader (Tecan, Männedorf, Switzerland). 96-well polystyrene microplates (Brand, Wertheim, Germany) were coated with poly-L-lysine. 30.000 cells per well were seeded during transfection 48h prior the measurement. Before the measurement, the culture medium was removed, and the cells were washed with external buffer. Subsequently, 80 µl of external buffer containing 1 µM 6H-F-Colenterazine ^8^ was added into each well and incubated for 15 minutes. The measurements were conducted at 37°C. For the recording of dose-response curves the following protocol was used: first, the baseline-BRET was recorded. After ten cycles, 20 µl of agonist containing solution or external buffer, as negative control, was added (agonist phase). After a further ten cycles, 20 µl of the saturating concentration of the respective agonist or external buffer was added, and 10 more cycles were measured (saturation phase). The measurements were controlled using the software SparkControlTM. The donor luminescence (nLuc) was recorded at 415 to 470 nm and acceptor luminescence (YFP) was recorded at 520 to 590 nm, respectively, and the (nLuc/CFP) luminescence ratio was calculated. The negative control trace with just of external buffer was subtracted and the trace was normalized to the baseline luminescence. For the concentration-response curve, the mean of the respective agonist phase was normalized to the saturation phase. Each n indicates one transfection measured as triplicate.

### Expression controls via Flow Cytometry

Surface expression of the chimeric constructs was analyzed using flow cytometry (Supplemental Figure 2p-r, 4p-r). For this, cells were transfected as described above and seeded to 24-well plates. After 48h, cells were washed with PBS and detached with trypsin, and resuspended with DMEM. The harvested cells were centrifuged at 2000 rpm for 5 min, washed with PBS and again centrifuged. The muscarinic mCherry-receptor constructs were analyzed via flow cytometry using a Guava easyCyte 6-2L system flow cytometer (Merck Millipore, Darmstadt, Germany). Every experiment was performed with 5000 cells out of minimum three independent transfections in two to three replicates each condition. Data was analyzed using the GuavaSoft Software 4.5 package.

### Data analysis and Statistics

Single-cell FRET-measurements were corrected for background fluorescence, bleed-through of CFP into YFP channel and false excitation of YFP using Microsoft Excel and were corrected for photobleaching (using OriginPro2024). BRET-measurements were corrected for background luminescence using Microsoft Excel and further data analysis was performed using GraphPad Prism 10 (GraphPad Software). Data is shown as mean ± SEM and group size defined as n. Statistical analyses were performed with an ordinary one-way ANOVA with Dunnet’s T3 multiple comparisons test, as indicated. Differences were considered as statistically significant if p ≤ 0.05. Concentration-response curves were fitted with a non-linear least-squares fit with variable slope and a constrained top and bottom using the following function:

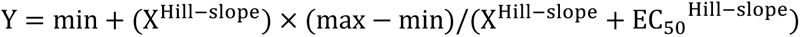

where min and max are the minimal and maximal response and EC_50_ is the half-maximal effective concentration. Stability of the receptor – G protein interactions was analyzed as described before ^5^. For this, the baseline was subtracted to 0 at the level of GTPγS application and the FRET ratio Δ(F_YFP_/F_CFP_) was normalized to the second agonist peak and set to 1. The stability of the interaction was quantified by the AUC (calculated by the trapezoidal method in Microsoft Excel) to compare the different types of dissociation kinetics.

## Author contributions

S.B.K., V.J., K.K. and A.L.K. performed experiments, S.B.K. analyzed the data and wrote the manuscript, M.B. supervised the study and edited the manuscript.

## Competing interests

The authors declare no competing interests.

**Supplemental Information** is available for this paper.

## Notes

### Competing Interest Statement

The authors have declared no competing interest.

