## Supplemental information for "Dissecting GPCR Selectivity: A complex interplay of various intracellular motifs determines G-protein binding and activation"

**Affiliations:**

Table 1: Chimeric Receptor Constructs

| Construct | Position removed from WT Receptor | Position inserted from other receptor |
|---|---|---|
| M3 ICL2 M2 | R177-A178 | P312-V133 |
| M3 ICL3 M2 | E274 - T471 | D299 - K367 |
| M3 TM56 M2 | Y251-E274 & T471-A495 | Y206-D299 & K367-A391 |
| M3 TM56 + H8 M2 | Y251-E274 & T471-A495 & K549- D564 | Y206-D299 & K367-A391 & A445- Y459 |
| M3 TM56 + PBC M2 | Y251-E274 & T471-A495 & K565-K570 | Y206-D299 & K367-A391 & K460-R466 |
| M3 ICL3 + TM56 M2 | Y251- A495 | Y206-A391 |
| M3 ICL3+ TM56 + PBC M2 | Y251- A495 & K565-K570 | Y206-A391 & K460-R466 |
| M3 H8 M2 | K549-D564 | A445-Y459 |
| M3 H8 + CTerm M2 | K549-L590 | A445-A466 |
| M3 PBC M2 | K565-K570 | K460-R466 |
| M3 PBC short | K565- L590 | K460-R466 |
| M3 -distal Cterm | Q571-L590 | - |
| M3 ICL2 + ICL3 + TM56 M2 | R177-A178 & Y251- A495 | P312-V133 & Y206-A391 |
| M3 complete M2 | R177-A178 & Y251- A495 & K549-L590 | P312-V133 & Y206-A391 & A445-A466 |
| M2 ICL2 M3 | P312-V133 | R177-A178 |
| M2 ICL3 M3 | D299 - K367 | E274 - T471 |
| M2 TM56 M3 | Y206-D299 & K367-A391 | Y251-E274 & T471-A495 |
| M2 TM56 + H8 M3 | Y206-D299 & K367-A391 & A445- Y459 | Y251-E274 & T471-A495 & K549- D564 |
| M2 TM56 + PBC M3 | Y206-D299 & K367-A391 & K460-R466 | Y251-E274 & T471-A495 & K565-K570 |
| M2 ICL3 + TM56 M3 | Y206-A391 | Y251- A495 |
| M2 ICL3+ TM56 + PBC M3 | Y206-A391 & K460-R466 | Y251- A495 & K565-K570 |
| M2 H8 M3 | A445-Y459 | K549-D564 |
| M2 H8 + CTerm M3 | A445-A466 | K549-L590 |
| M2 ICL3 + TM56 + H8 + Cterm | Y206-A391 & A445-A466 | Y251- A495 & K549-L590 |
| M2 PBC M3 | K460-R466 | K565-K570 |
| M2 +distal Cterm M3 | WT | Q571-L590 added |
| M2 ICL2 + ICL3 + TM56 M3 | P312-V133 & Y206-A391 | R177-A178 & Y251- A495 |
| M2 complete M3 | P312-V133 & Y206-A391 & A445-A466 | R177-A178 & Y251- A495 & K549-L590 |

Table 2: Oligonucleotide Primers

| Construct | Primer Sequence 5' → 3' |
|---|---|
| mCherry-M3 | AAA AAA GAA TTC CTT GTA CAG CTC GTC CAT GCC<br>AAAAAA GAA TTC ATG ACC TTG CAC AAT AAC AGT ACA ACC<br>CGCCCTGAGCTACATCTTCTGCCTGGTATTCGCCATGGTGAGCAAGGGCGAG<br>AGAAGATGTAGCTCAGGGCGATGATCGTCTTCATGGATCCGAGCTCGGTACC<br>GCAGGCCTTGTAGCGGCCGCTCGAG<br>GCAAGGTCATGAATTCCTTGTACAGCTCGTCCATGCC<br>CAAGGAATTCATGACCTTGCACAATAACAGTACAACC<br>GCGGCCGCTACAAGGCCTGCTCGGGTG<br>AGGCCTTGTAGCGGCCGCTCGAGGGGGGGC |
| M2-mCit | Digestion and Ligation with XbaI and NotI from M3-mCit |
| mCherry-M2 | AAA AAA GAA TTC ACC ATG AAG ACG ATC ATC GCC<br>AAA AAA GAA TTC CTT GTA CAG CTC GTC CAT GCC |
| PTX-A pcDNA3 | Fw: AAAAA AAGCTT ATGGACGATCCTCCCGC<br>Rev: AAAAA TCTAGA ACGAATACGCGATGCTTTC |
| **M3 Chimeric Receptors** | |
| M3 ICL2 M2 | GGCCGCTCACGTACC**C**AG**T**CAAACGAACAACAAAG |
| M3 ICL3 M2 | BB fw: CTAAGCAGCCTGCAAAAAAGAGGATGTCCCTGGTCAAGGAG<br>BB rev: GGGTCTTGGTTGGCAACAGGCAGGCCAGCAAGCTC<br>Insert fw: CCAAAGAGCTTGCTGGCCTGCCTGTTGCCAACCAAGACC<br>Insert rev: TCCTTGACCAGGGACATCCTCTTTTTTGCAGGCTGCTTAGTCATC |
| M3 TM56 M2 | Same as M2 ICL3 M3, Template ICL3 + TM56 M2 |
| M3 TM56 + H8 M2 | Same as M3 H8 M2, Template M3 TM56 M2 |
| M3 TM56 + PBC M2 | Same as M3 pbc M2, Template M3 TM56 M2 |
| M3 ICL3 + TM56 M2 | BB fw: CCA AGA GGT TTG CTC TGA AGA AGA TGA CTA AGC AGC CTG CAA AAA AG<br>BB rev: ACA AAG TTT TCT GTC TCT GCG TCT TGG TTG GCA ACA GGC T<br>Insert fw: AGC CTG TTG CCA ACC AAG ACG CAG AGA CAG AAA ACT TTG TCC ACC<br>Insert rev: GCA GGC TGC TTA GTC ATC TTC TTC AGA GCA AAC CTC TTG GCC |
| M3 ICL3+ TM56 + PBC M2 | Same as M3 pbc M2, Template M3 ICL3 + TM56 M2 |
| M3 H8 M2 | Fw: acaccttctcatgtgtcattatAAAAAAAAGAGGCGCAAG<br>Rev: ttaaaggtcttcttgaaggtggcGTTGCACAGAGCATAGCA |
| M3 H8 + CTerm M2 | BB fw: TACGGCAAGCTGACCCTGAAG<br>BB rev: TTGAAGGTGGCGTTGCACAGAGCATAGCACACG<br>Insert fw: CTGTGCAACGCCACCTTCAAGAAGACCTTTAAACAC |

| | |
|---|---|
| | Insert rev: CTTCAGGGTCAGCTTGCCGTAG |
| M3 PBC M2 | Fw: gcgctacaaggCAGCAGTACCAGCAG<br>Rev: ctatgttcttGTCACACTGGCACAG |
| M3 PBC short | Fw: TCTAGAGTGAGCAAGGGC rev: CCTTGTAGCGCCTATGTTC |
| M3 -distal Cterm | Fw: TCTAGAGTGAGCAAGGGC rev: CTTGCGCCTCTTTTTTTTG |
| M3 ICL2 + ICL3 + TM56 M2 | Same as M3 ICL2 M2, Template M3 ICL3 + TM56 M2 |
| M3 complete M2 | Template M2 ICL2 + ICL3 + TM56 M2<br>Insert Fw: TGTGCTATGCTCTGTGCAACGCCACCTTCAAGAAGACCTTTAAACAC<br>Insert rev mCit: TCGCCCTTGCTCACTCTAGACCTTGTAGCGCCTATGTTCTTATAATGACAC<br>Insert rev mCherry: CCCCCTCGAGCGGCCGCTACCTTGTAGCGCCTATGTTCTTATAATGACAC<br>BB Rev: AAGGTCTTCTTGAAGGTGGCGTTGCACAGAGCATAGCACACG<br>BB fw mCit: AGAACATAGGCGCTACAAGGTCTAGAGTGAGCAAGGGCGA<br>BB fw mCherry: AGAACATAGGCGCTACAAGGTAGCGGCCGCTCGAGG |
| **M2 Chimeric Recpetors** | |
| M2 ICL2 M3 | CAAAACCTCTGACCTACC**G**AG**C**CAAGCGGACCACAAAAATG |
| M2 ICL3 M3 | BB fw: GTCAGATCACTAAGCGGAAAAAGCCTCCTCCTTCCCG<br>BB rev: GCCTCTGTCCCAGAGGCTTGCTCCTTCTTGTCCTTCTTTATCCTG<br>Insert fw: TAAAGAAGGACAAGAAGGAGCAAGCCTCTGGGACAGAG<br>Insert rev: TCCCGGGAAGGAGGAGGCTTTTTCCGCTTAGTGATCTGACTTCTGG |
| M2 TM56 M3 | BB fw: TTG TAG CCC GCA AGA TTG TGA CCA GAA GTC AGA TCA CTA AGC GG<br>BB rev: ACC AGA CTT GGA GAA ACG GGC TCT GTC CCA GAG GCT TGC A<br>Insert fw: TGC AAG CCT CTG GGA CAG AGC CCG TTT CTC CAA GTC TGG TAC A<br>Insert rev: TTA GTG ATC TGA CTT CTG GTC ACA ATC TTG CGG GCT ACA ATA TTC T |
| M2 TM56 + H8 M3 | Same as M2 H8 M3, Template M2 TM56 M3 |
| M2 TM56 + PBC M3 | Same as M2 PBC M3, Template M2 TM56 M3 |
| M2 ICL3 + TM56 M3 | Insert fw: GTG CGA TTC TGT TGG CTT TCA TCA TCA CTT GG<br>Insert rev: TAG ATC CTC CAG TAT AGC ACA GTC ATG ATG ATC ACT GGC<br>BB fw: TGT GCT ATA CTG GAG GAT CTA TAA GGA AAC TGA AAA GCG<br>BB rev: GAA AGC CAA CAG AAT CGC ACT GAG GGT CTG GG |
| M2 ICL3+ TM56 + PBC M3 | Template: M2 PBC M3<br>BB fw: GCCCAGACCCTCAGTGCGATTCTGTTGGCTTTCATCATCACTTGGG<br>BB rev: CTTATAGATCCTCCAGTATAGCACAGTCATGATGATCACT<br>Insert fw: AGTGATCATCATGACTGTGCTATACTGGAGGATCTATAAGGAAACTGAAAAGCG<br>Insert rev: TGATGATGAAAGCCAACAGAATCGCACTGAGGGTCTGGG |

| | |
|---|---|
| M2 H8 M3 | atgctgctgctgtgccagtgtgacAAGAACATAGGCGCTACAAG<br>cttgaaagtggttctgaatgttttATTGCAAAGTGCATAGCAG |
| M2 H8 + CTerm M3 | BB fw: TACGGCAAGCTGACCCTGAAG<br>Insert rev: CTTCAGGGTCAGCTTGCCGTAG |
| M2 ICL3+TM56 + H8 + Cterm | Same as M2 ICL3+TM56 M3, Template M2 H8 + CTerm |
| M2 PBC M3 | mCit: TGACGGCCGCTCGAG; CTTGCGCCTCTTTTTTTTATAATGACAC<br>mCherry: TCTAGAGTGAGCAAGG; CTTGCGCCTCTTTTTTTTATAATG |
| M2 +distal Cterm M3 | mCherry: fw: tttcacaagcgcgcacccgagcaggccttgTGAGCGGCCGCTCGAGCA<br>rev: aatgaccgactgtctctgctggtactgctgCCTTGTAGCGCCTATGTTCTTATAATGACACATG |
| M2 ICL2 + ICL3 + TM56 M3 | Same as M2 ICL2 M3, Template M2 ICL3 + TM56 M3 |
| M2 complete M3 | Template M2 ICL2 + ICL3 + TM56<br>BB fw mCherry: TTTCACAAGCGCGCACCCGAGCAGGCCTTGTGATCGAGGGGGGGCCCTATTC<br>BB fw mCit: AGAACATAGGCGCTACAAGGTCTAGAGTGAGCAAGGGCGAG<br>BB rev: AAAGTGGTTCTGAATGTTTTATTGCAAAGTGCATAGCAGGCA<br>Insert fw: CCTGCTATGCACTTTGCAATAAAACATTCAGAACCACTTTCAAGATGCT<br>Insert rev mCherry: ATAGGGCCCCCCCTCGATCACAAGGCCTGCTCGGGTG<br>Insert rev mCit: TCGCCCTTGCTCACTCTAGACCTTGTAGCGCCTATGTTCTTATAATGACAC |

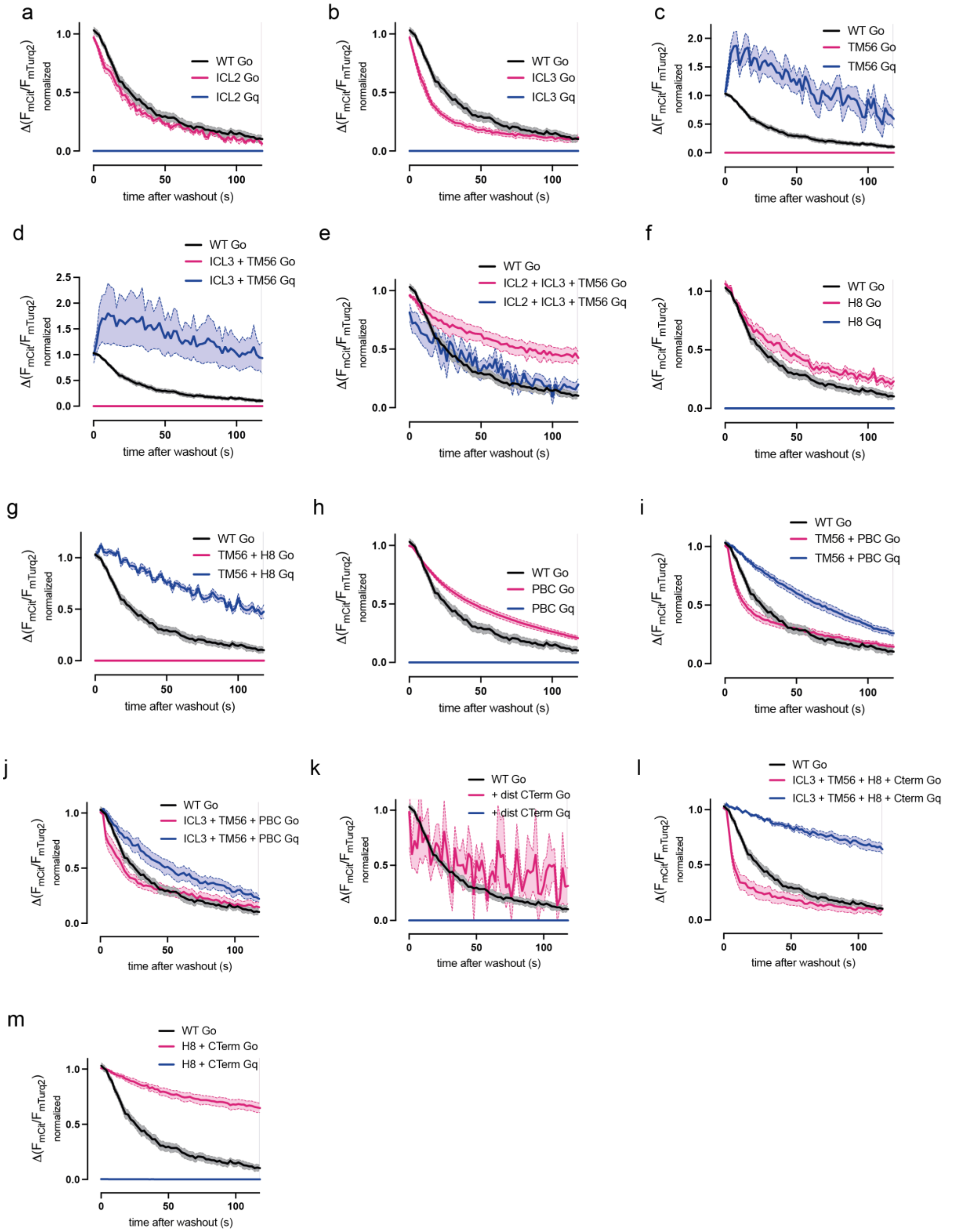


***Supplemental Figure 1: M2-based chimeric receptors enable the binding of Go and Gq proteins by the M2 receptor. a-m*** *Averaged FRET-based measurement of M2 receptor-G-protein interaction in transfected, permeabilized HEK293T cells expressing a mCit-tagged receptor-chimera and the respective G-proteins with a Gγ carrying a mTurq with different chimeric receptors. Wash-out of the second 10 µM acetylcholine application*

by buffer solution is depicted (measurement protocol as described in Figure 1c). The area under the curve was calculated to quantify the strength of the interaction and plotted as bar graph In Figure 2 and 4a-b (Mean ± SEM, n=7-13).

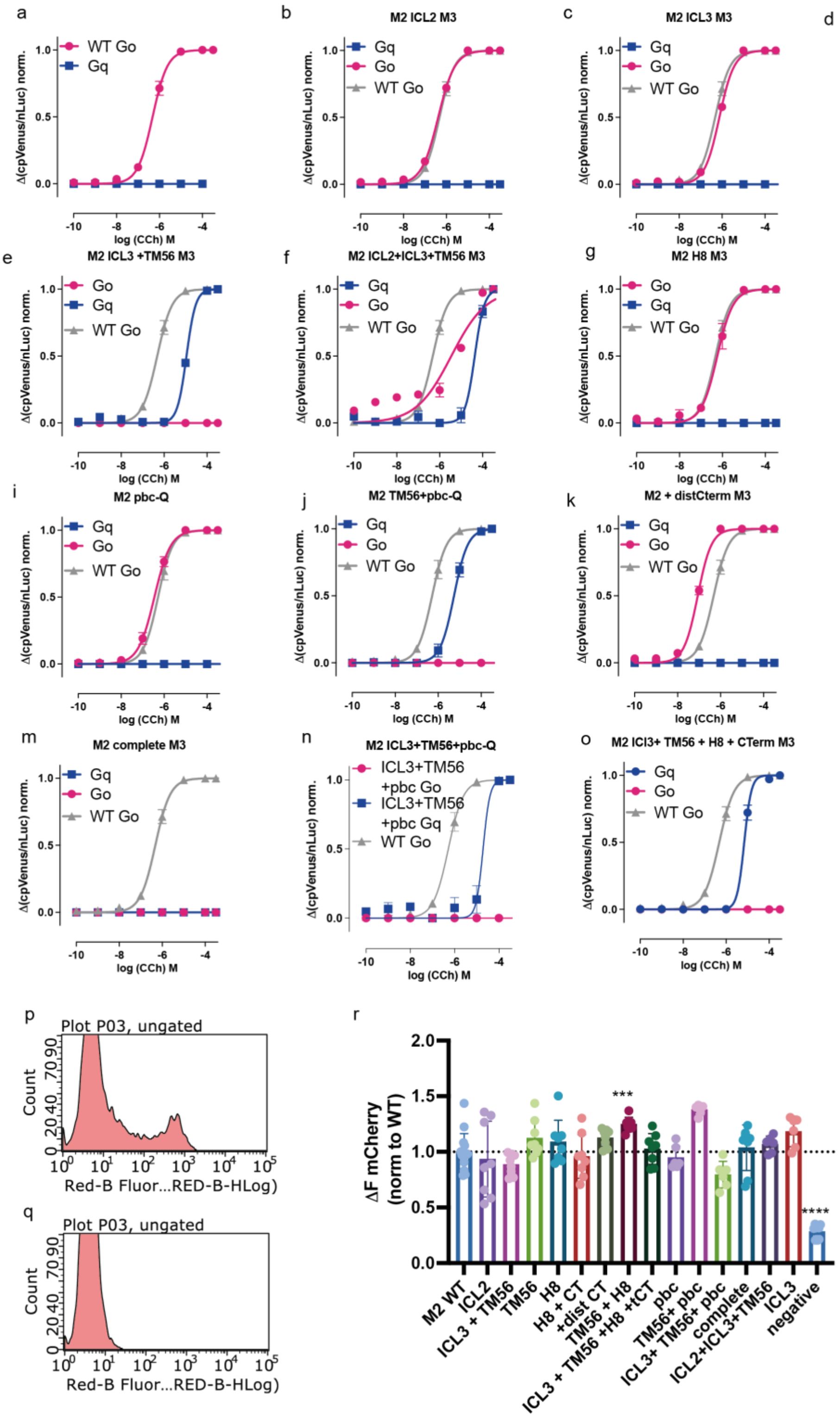

***Supplemental Figure 2: M2-based chimeric receptors enable the subsequent activation of Go Gq proteins by the M2 receptor. a-o*** *Concentration-response curves for carbachol induced G-protein activation, measured using the BRET-based G-case assays in multiwell format in a plate reader. HEK293T cells were transfected with the G-protein activity sensor and the chimeric M2-based receptor carrying an N-terminal fused mCherry. Concentration-response curves were fitted and the calculated EC50-values were plotted in bar graphs (Figure 2 and 4c-d). Measurements were performed in triplicates for each transfection, with at least three independent transfections.* ***p-r*** *Expression control of chimeric M2-based receptors using flow cytometry. HEK293T cells transfected with N-terminal mCherry tagged WT receptor (p), chimeric receptors or no receptor (q) were analyzed via flow cytometry regarding their red fluorescence (p-q). The mean fluorescence was normalized to the fluorescence of the WT receptor and plotted as bar graph (r). Measurements were performed in triplicates for each transfection, with at least three independent transfections. (Statistic: Ordinary One-way ANOVA with Dunnett's multiple comparison test, **** p<0.0001, *** p<0.001, ns: p>0.05).*

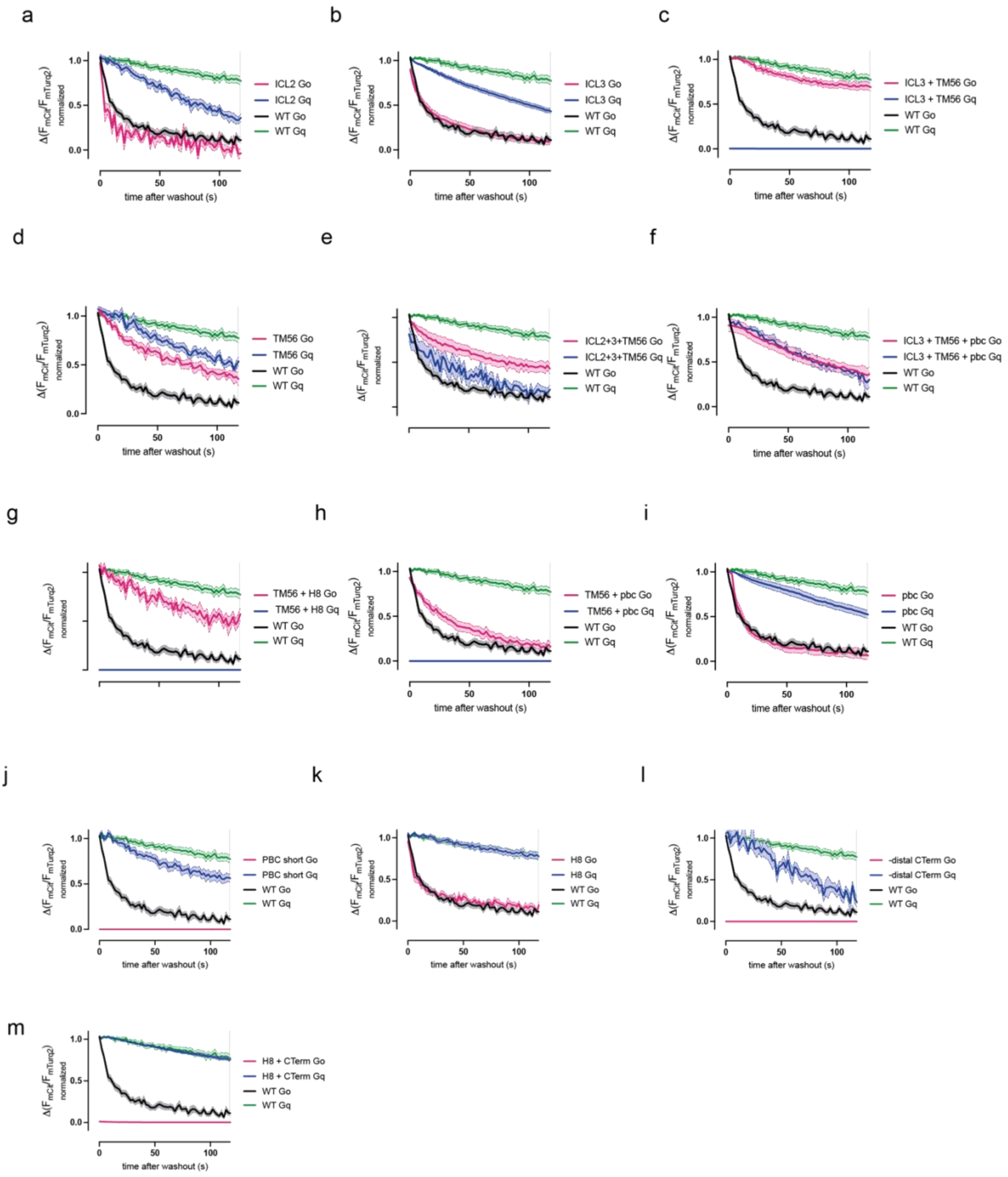


***Supplemental Figure 3: M3-based chimeric receptors can be modified to signal solely via Gq or Go. a-m*** *Averaged FRET-based measurement of M3 receptor-G-protein interaction in transfected, permeabilized HEK293T cells expressing a mCit-tagged receptor-chimera and the respective G-proteins with a G$\gamma$ carrying a mTurq with different chimeric receptors. Wash-out of the second 10 µM acetylcholine application by buffer solution is depicted (measurement protocol as described in Figure 1c). The area*

*under the curve was calculated to quantify the strength of the interaction and plotted as bar graph In Figure 3 and 4e-f (Mean ± SEM, n=10-14).*

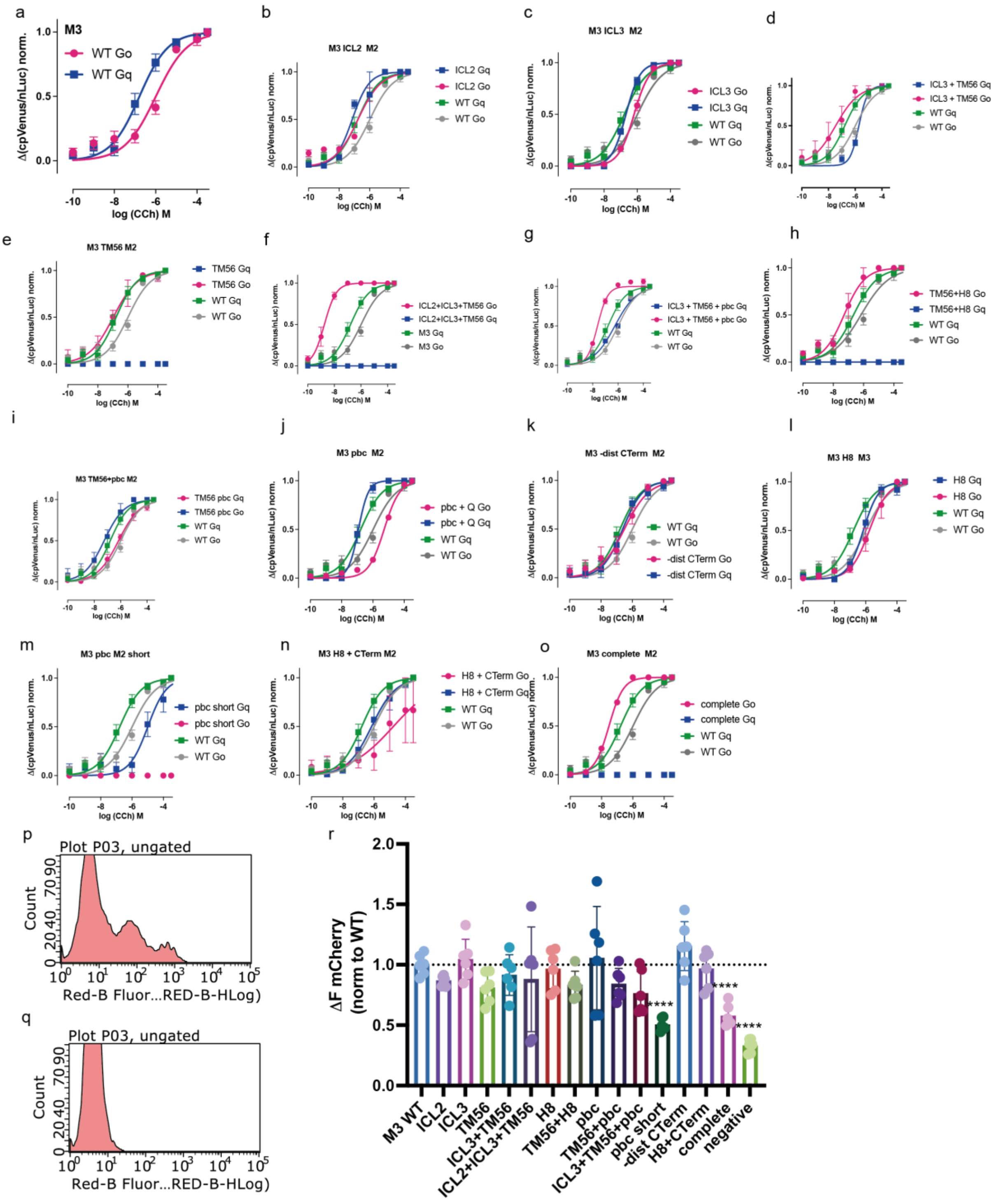


***Supplemental Figure 4: M3-based chimeric receptors can be modified to signal solely via Gq or Go. a-o*** *Concentration-response curves for carbachol induced G-protein activation, measured using the BRET-based G-case assays in multiwell format in a plate reader. HEK293T cells were transfected with the G-protein activity sensor and the chimeric M3-based receptor carrying an N-terminal fused mCherry. Concentration-response curves were fitted and the calculated EC50-values were plotted in bar graphs*

*(Figure 3 and 4g-h). Measurements were performed in triplicate for each transfection, with at least three independent transfections.* ***p-r*** *Expression control of chimeric M3-based receptors using flow cytometry. HEK293T cells transfected with N-terminal mCherry tagged WT receptor (p), chimeric receptors or no receptor (q) were analyzed via flow cytometry regarding their red fluorescence (p-q). The mean fluorescence was normalized to the fluorescence of the WT receptor and plotted as bar graph (r). Measurements were performed in triplicates for each transfection, with at least three independent transfections. (Statistic: Ordinary One-way ANOVA with Dunnett's multiple comparison test, ***** $p<0.0001$*, ns:* $p>0.05$*).*
